# Activation and desensitization mechanism of AMPA receptor-TARP complex by cryo-EM

**DOI:** 10.1101/158402

**Authors:** Shanshuang Chen, Yan Zhao, Yuhang (Steven) Wang, Mrinal Shekhar, Emad Tajkhorshid, Eric Gouaux

## Summary

AMPA receptors mediate the majority of fast excitatory neurotransmission in the mammalian brain and transduce the binding of presynaptically released glutamate to the opening of a transmembrane cation channel. Within the postsynaptic density, however, AMPA receptors coassemble with transmembrane AMPA receptor regulatory proteins (TARPs), yielding altered gating kinetics, receptor pharmacology and pore properties. Here we elucidate full-length GluA2-TARP γ2 complex structures in the presence of the partial agonist kainate or the full agonist quisqualate together with a positive allosteric modulator, and with quisqualate alone. We show how TARPs sculpt the ligand binding domain gating ring, enhancing kainate potency and diminishing the ensemble of desensitized states. The 4 TARPs encircle receptor ion channel, stabilizing M2 helices and pore loops, thus showing how TARPs alter receptor pore properties. Structural and computational analysis suggests the full agonist/modulator complex harbors an ion-permeable channel gate, thus providing the first view of an activated AMPA receptor.

In the mammalian central nervous system the majority of fast excitatory neurotransmission is initiated by α-amino-3-hydroxy-5-methyl-4-isoxazole propionic acid (AMPA)-sensitive ionotropic glutamate receptors in complex with modulatory auxiliary subunits (Jackson and Nicoll, 2011; Traynelis et al., 2010). Transmembrane AMPA receptor regulatory proteins (TARPs) (Chen et al., 2000), the most widespread and well studied family of auxiliary proteins, alter AMPA receptor gating kinetics, ion channel properties and pharmacology (Milstein and Nicoll, 2008). The prototypical TARP, deemed stargazin or TARP γ2, potentiates AMPA activity by decelerating deactivation and desensitization kinetics, facilitating recovery from desensitization, boosting the efficacy of partial agonists, and attenuating polyamine block (Milstein et al., 2007; Soto et al., 2007; Tomita et al., 2005).

AMPA receptors have a modular architecture with synaptically localized amino terminal domains (ATDs) and ligand binding domains (LBDs), an ion channel forming transmembrane domain (TMD) and a largely unstructured cytoplasmic domain (CTD) (O’Hara et al., 1993; Soderling and Derkach, 2000; Stern-Bach et al., 1994; Wo and Oswald, 1995; Wollmuth and Sobolevsky, 2004). Extensive studies on isolated receptor domains and intact receptors have illuminated, at high resolution, how agonists induce local ‘clamshell’ closure of the LBDs and how the LBDs are arranged as nonequivalent pairs of A/C and B/D dimers within an overall 2-fold symmetric LBD ‘gating ring’ (Armstrong and Gouaux, 2000; Jin et al., 2009; Kuusinen et al., 1999; Sobolevsky et al., 2009; Sun et al., 2002). Although crystallographic and cryo-EM structures of intact receptors have been determined in the presence of partial and full agonists (Chen et al., 2014; Dürr et al., 2014; Meyerson et al., 2014), no studies have yet captured the ion channel gate in an open conformation. Indeed, the x-ray studies suggest that upon receptor activation, not only does the LBD gating ring expand but it also moves closer to TMD (Chen et al., 2014; Dürr et al., 2014). However, these structural studies were carried out on thermostabilized receptor variants with low open probabilities and they showed that the ion channel gate remained closed, thus suggesting that despite gating ring expansion, ‘compression’ of the LBD toward the membrane bilayer decoupled agonist-binding from ion channel gating.

Multiple studies on isolated domains and on the intact receptor have also provided insights into the structural underpinnings of receptor desensitization. Fortuitous cysteine mutagenesis and electrophysiological studies carried out on the intact and ΔATD receptor, along with crystallographic studies on the isolated LBD ‘dimers’ suggested that cleavage of the LBD dimer D1-D1 interface was sufficient to promote receptor desensitization (Armstrong et al., 2006). By contrast, cryo-EM studies of isolated AMPA receptors and the closely related kainate receptors suggest that there are, instead, large scale rearrangements of the LBD layer from 2-fold to ∼4-fold symmetry (Meyerson et al., 2016; Meyerson et al., 2014). Indeed, a low resolution x-ray study of the intact receptor, as well as cryo-EM studies, are also suggestive of large scale LBD rearrangements of the LBD layer upon receptor desensitization (Dürr et al., 2014).

At present, there are no structural studies of an AMPA receptor in an activated state with an open ion channel gate, nor are there studies of the AMPA receptor-TARP complex in multiple ligand-bound conformations. Moreover, there are no structural insights into the conformational ensemble of structures associated with an AMPA receptor – TARP complex upon receptor desensitization. To gain insight into how TARP subunits modulate receptor activity, from increasing the efficacy of partial agonists to altering the properties of the ion channel pore, we carried out cryo-EM reconstructions on the full length GluA2 AMPA receptor in complex with intact TARP γ2 auxiliary subunits. These studies not only show how TARP subunits modulate receptor activation and recovery from desensitization, but they also provide the first view of an AMPA receptor TMD, thus lending new insight into gating, permeation and block of AMPA receptors.

## Structure determination

To elucidate the molecular mechanism for partial and full agonist action on AMPA receptors bound to TARP subunits we determined cryo-EM structures of the GluA2-TARP γ2 complex with the classic partial agonist kainate (Patneau et al., 1993) and the high affinity full agonist quisqualate (Jin et al., 2002), in the presence of the potent, 2-fold symmetric positive allosteric modulator (R,R)-2b (Figures 1A-1C) (Kaae et al., 2007). The three-dimensional (3D) classification of the quisqualate/(R,R)-2b complex revealed four classes featuring ∼4-fold related protrusions on the extracellular side of the detergent micelle, consistent with four TARP γ2 subunits encircling the receptor TMD (Figure S1). The two remaining classes lacked prominent protrusions from the detergent micelle and thus are likely composed of particles not saturated with TARP subunits. Hence, we excluded them from further analysis (Figure S1). Inspection of the four classes with TARP-like protrusions indicated that the most prominent differences between each class involved the conformationally mobile ATD layer, and further 3D-classification on the combined four classes focused on the LBD-TMD layer did not suggest evidence of discernable conformational heterogeneity (Figure S1).

**Figure 1.**
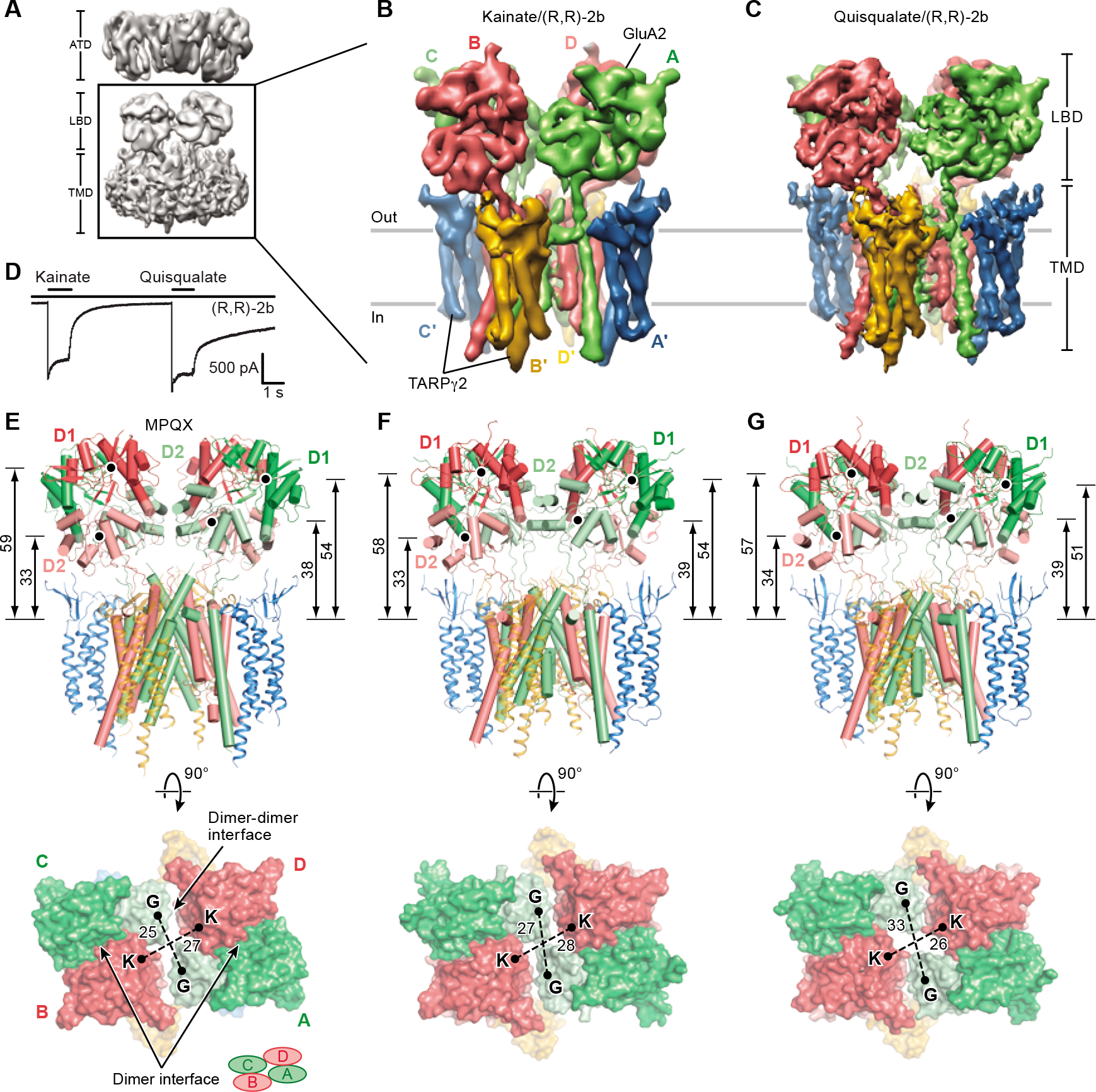
Activated states of the GluA2-TARP γ2 complex. (A) Initial cryo-EM reconstruction of full-length GluA2-TARP γ2 complex bound with quisqualate and (R,R)-2b. Higher resolution reconstructions focused on the LBD/TMD layers. (B) and (C) Cryo-EM reconstructions of GluA2-TARP γ2 complex bound with kainate and quisqualate in the presence of (R,R)-2b, respectively. (D)-(F) Structures of the LBD/TMD layers in the previously determined MPQX complex (Zhao et al., 2016) (D), with kainate/(R,R)-2b (E) and with quisqualate/(R,R)-2b (F). The LBD D1 and D2 lobes are shown in darker and lighter shades, respectively. COMs of D1 and D2 are indicated by black dots, whose distances from a reference point, COM of Thr617 Cα atoms, are labeled in the “side views”; COM distances between proximal helices between opposing subunits, helix G (G) for A/C subunits and helix K (K) for B/D subunits, are indicated in the “top-down views”. Distances are in angstroms (Å).

We thus focused our efforts on the domains most important to receptor activation - the LBD and TMD layers - and excluded the conformationally mobile ATD layer by way of a soft mask, carrying out 3D-reconstruction of the GluA2-TARP γ2 complex (Figure S1). The overall resolutions estimated for the cryo-EM density maps of the GluA2-TARP γ2 complex bound with quisqualate/(R,R)-2b and kainate/(R,R)-2b were 4.9 Å and 6.4 Å, respectively (Figures 1A-1C and S2A-S2J). Both maps share similar overall features. The LBD subunits are organized as 2-fold symmetric dimers-of-dimers ‘above’ an approximately 4-fold symmetric TMD that, in turn, is surrounded by a ∼4-fold symmetric ensemble of TARP subunits (Figures 1B and 1C). In harmony with previous structures of AMPA receptors in non desensitized states, there are two conformationally distinct pairs of receptor subunits, deemed A/C and B/D, along with 4 TARP subunits defined as A’/C’ and B’/D’ (Figures 1B and 1C).

Structural modeling of the quisqualate/(R,R)-2b bound GluA2-TARP γ2 complex was carried out by rigid body fitting of LBD, TMD and TARP domains extracted from known structures into the density map. This resulted in an initial model composed of separately docked D1 and D2 lobes of the LBD derived from the crystal structure of an isolated GluA2 LBD quisqualate complex (Jin et al., 2002), receptor TMD from the crystal structure of an intact GluA2 receptor (Sobolevsky et al., 2009) and TARPs from the single particle cryo-EM structure of the GluA2-TARP γ2 complex with MPQX (Zhao et al., 2016). Manual adjustment of secondary structure elements was carried out where merited by the quality of the EM density map. The amino acid register throughout the receptor TMD was confirmed by well defined side-chain densities for aromatic residues (Figures S3A and S3B). However, the density map provided only continuous main-chain features for the S1-preM1 linkers, the M3-S2 linkers, the TARP TM2-TM3 linkers and the acidic loop adjacent to the α1 helix of the B’/D’ TARP subunits. Structural elements were not built for either the S2-M4 linkers or the M1-M2 linkers due to weak density (Figure S3A). The structural model was further improved by molecular dynamics flexible fitting (MDFF) (Trabuco et al., 2008), real-space refinement and manual adjustment, yielding a structure that correlates well with the EM map and bears excellent stereochemistry (Table S1). Because the density maps show little conformational difference in the TMD regions between the quisqualate/(R,R)-2b and kainate/(R,R)-2b complexes, a model for the kainate/(R,R)-2b bound GluA2-TARP γ2 was generated by fitting the receptor TMD and TARPs from the quisqualate/(R,R)-2b complex structure as a rigid body. The D1 and D2 lobes of the LBD were extracted from the crystal structure of the isolated LBD bound with kainate/(R,R)-2b and separately fit into the density map (Figure S3C) (Dürr et al., 2014), followed by refinement in real-space.

## Agonists expand LBD gating ring

Partial and full agonists elicit a progressive expansion of the LBD gating ring relative to the MPQX bound state (Twomey et al., 2016; Zhao et al., 2016), and the nonequivalent A/C and B/D LBDs adopt different positions and orientations relative to the TMD (Figures 1D-1F). As in the MPQX complex, LBD dimerization is mediated by extensive D1-D1 interactions and contacts with (R,R)-2b, independent of the degree of LBD clamshell closure. There is a less extensive dimer-dimer interface formed between proximal subunits in the quisqualate-activated complex or between both proximal and opposing subunits in the kainate-activated complex (Figures 1D-1F). To measure the changes in gating ring conformation, we used the centers of masses (COMs) of proximal helices from opposing subunits, helices G for A/C LBDs and helices K for B/D LBDs. These helices extend by 6 Å and contract by 2 Å, respectively, upon progression from the partial to full agonist bound states, illuminating how agonist efficacy and clamshell closure is translated into structural arrangement of the LBD gating ring (Figures 1D-1F and Movie S1).

## Activation by full and partial agonists

On the one hand, prior studies on the isolated LBD have established a correlation between agonist efficacy and the extent of LBD clamshell closure stabilized by agonist binding (Armstrong and Gouaux, 2000). On the other hand, several studies suggest that antagonists and partial agonists can give rise to fully closed LBD clamshells and that the difference between antagonists, partial agonists and full agonists is the probability of the LBD occupying the fully closed conformation (Ahmed et al., 2013; Ahmed et al., 2011). Because we now have a full length receptor/TARP assembly, devoid of thermostabilizing mutations, in complex with partial and full agonists, we can address this question directly. Using the D1 lobe as a reference, superposition of crystal structures of the isolated LBD clamshells bound with the agonist/(R,R)-2b on the receptor/TARP complexes described here results in nearly superimposable D2 lobes (Figure S3D), demonstrating largely unaltered LBD clamshell closure in the context of TARP-associated intact receptors. Interestingly, the degree of LBD clamshell closure differs by ∼1° between the A/C and B/D subunits of the kainate/(R,R)-2b bound complex, where there is 15° and 14° closure at the A/C and B/D positions, respectively, in comparison to the MPQX-bound structure (Figures 2A and 2B). The 25° degree of closure, also relative to the MPQX structure, is the same in all subunits of the quisqualate/(R,R)-2b bound complex (Figures 2A and 2C). While the difference in clamshell closure between the A/C and B/D subunits for the kainate complex is subtle, we speculate that it is due to the differential ‘pulling’ force of the LBD clamshell on the A/C versus the B/D M3 helices.

**Figure 2.**
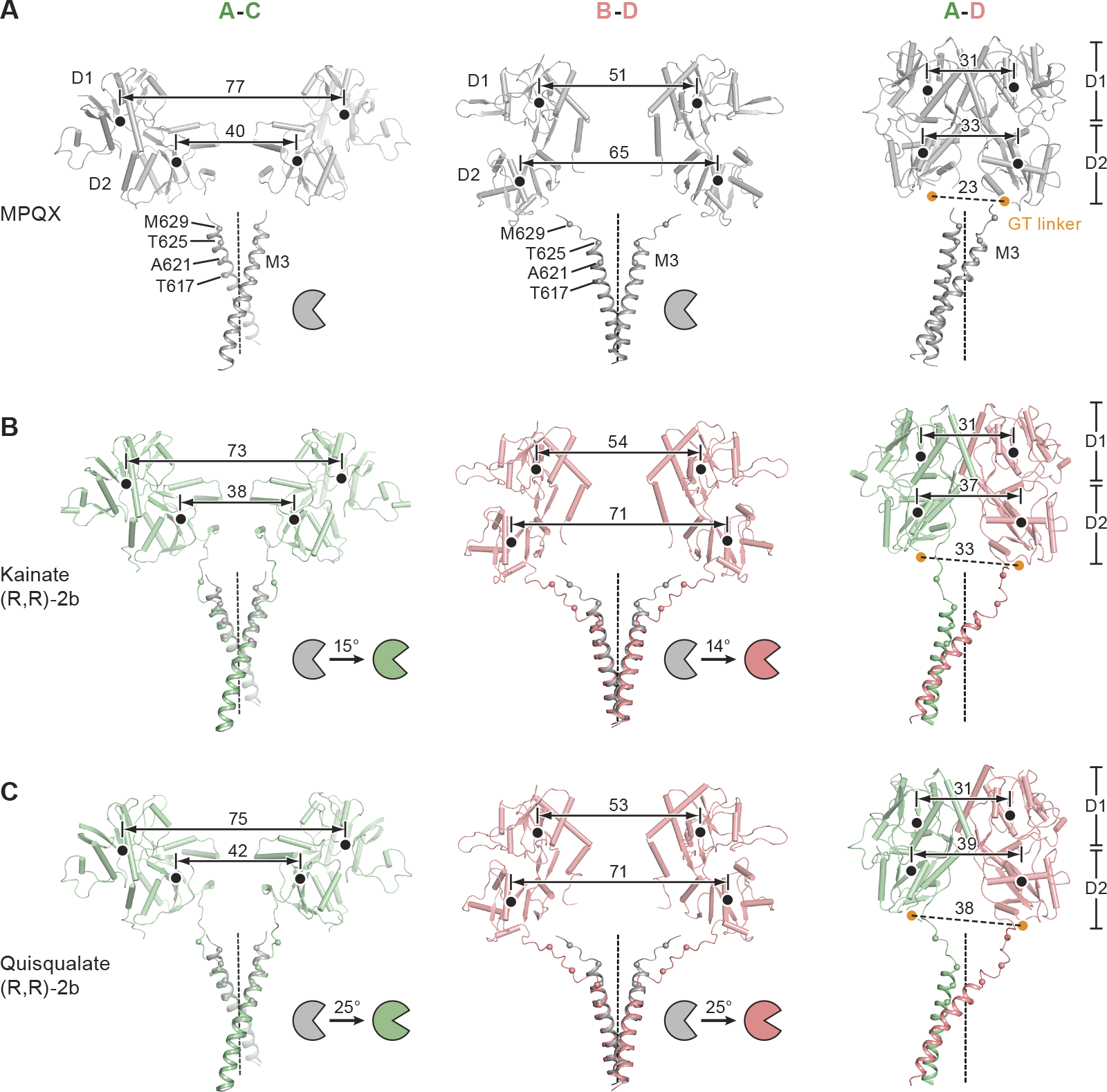
Conformational changes upon receptor activation. (A)-(C) Structural comparison of opposing subunit pairs, A/C and B/D, and adjacent subunits, A/D, showing the LBDs and M3 helices in the MPQX (A), kainate (B) and quisqualate (C) GluA2-TARP γ2 complexes. COMs of D1 and D2 lobes are indicated by black dots. Cα atoms of residues near M3 bundle crossing are shown as spheres. LBD clamshell closure is represented by schematic cartoons. Structures were superimposed using main-chain atoms of the M3 helices. In views showing LBD dimers (right panel), locations representing COMs of Gly-Thr linkers derived from isolated LBD structures are indicated by orange dots and were determined by superposition of a D2 lobe extracted from the crystal structure of an isolated LBD (PDB code: 1MM6) onto corresponding D2 lobes of the GluA2-TARP γ2 complexes. Distances are in angstroms (Å).

To elucidate how clamshell closure is transmitted to structural rearrangement of the channel gate, we compared the quisqualate-activated and kainate-activated GluA2-TARP γ2 complex structures with the MPQX complex (Figures 2B and 2C). Cognizant that the A/C and B/D LBDs occupy distinct positions in the receptor assembly, we see that agonist-induced clamshell closure causes only modest displacements in COMs of the A/C D2 lobes. By contrast, there is a 6 Å separation between COMs of the B/D D2 lobes, consistent with the B/D subunits exerting a greater ‘pulling force’ on the M3 helices (Figures 2B and 2C). Clear main-chain density indicates that the M3-S2 linkers at B/D positions in the quisqualate/(R,R)-2b and kainate/(R,R)-2b receptor-TARP complexes adopt a “coupled conformation” (Figures 2B and 2C) (Chen et al., 2014), where Ile633 is engaged within a hydrophobic pocket in the D2 lobe. Strikingly, the extracellular ends of the M3 helices, which include Met629, Thr625, Ala621 and Thr617, undergo deformation of helical secondary structure upon exertion of the pulling force transmitted from the LBD clamshell closure, via the M3-S2 linkers. At the A/C subunits Met629 Cα atoms undergo a nearly ‘vertical’ movement away from the membrane plane while at the B/D subunits the movement is in an orthogonal plane, nearly parallel with the membrane, showing that the A/C and B/D LBDs exert pulling forces in different directions (Figures 2B, 2C and Movie S1). In the context of the local LBD dimers, the D1-D1 LBD dimer interface is maintained during receptor-TARP activation by the modulator (R,R)-2b, and thus in the presence of either kainate or quisqualate, LBD clamshell closure is coupled to the separation of the D2 lobes and the M3-S2 linkers. More profound clamshell closure induced by the full-agonist quisqualate in comparison with the partial-agonist kainate yields larger D2-D2 separation (Figures 2B and 2C) quantified by distances between COMs of the D2 lobes and by positions equivalent to the “Gly-Thr” linker of isolated LBDs (Armstrong and Gouaux, 2000).

## Architecture of the ion channel pore

The density throughout the receptor TMD, including the M2 pore helix, the pore ‘loop’ and the canonical M3 gating helices of the quisqualate/(R,R)-2b bound complex is well defined, allowing us to reliably position main-chain and bulky side-chain groups, thus defining the most complete structure of an AMPA receptor ion channel pore to date (Figures 3A, S2E, S3A and S3B). This improved structural model for the pore region reveals key residues that stabilize the pore architecture and define ion channel properties. The M2 helices and the pore loop are largely positioned by interactions with the M1 and M3 helices within a subunit and from the Ml helix of an adjacent subunit, contacts mediated in part by aromatic residues resolved in the density map (Figure 3B). Trp605 (M3), Tyr533 (M1) and Trp605 from an adjacent Ml subunit form hydrophobic interactions with the C-terminal end of the M2 helix, whereas Phe541 (M1) stabilizes the *N*-terminus of the M2 helix (Figure 3B). These well defined interactions in the receptor TMD are in part due to the indirect interactions with TARP, via contacts between receptor M1 and M4 helices with TARP TM3 and TM4 elements. Indeed, superposition of the isolated receptor TMD with the receptor TMD from the TARP complex shows that the presence of TARP results in a large scale adjustment of receptor TMD interactions that not only allow for extensive receptor – TARP interactions but that also reduce the conformational mobility of the receptor M2 helix and pore loop (Figure S5A).

**Figure 3.**
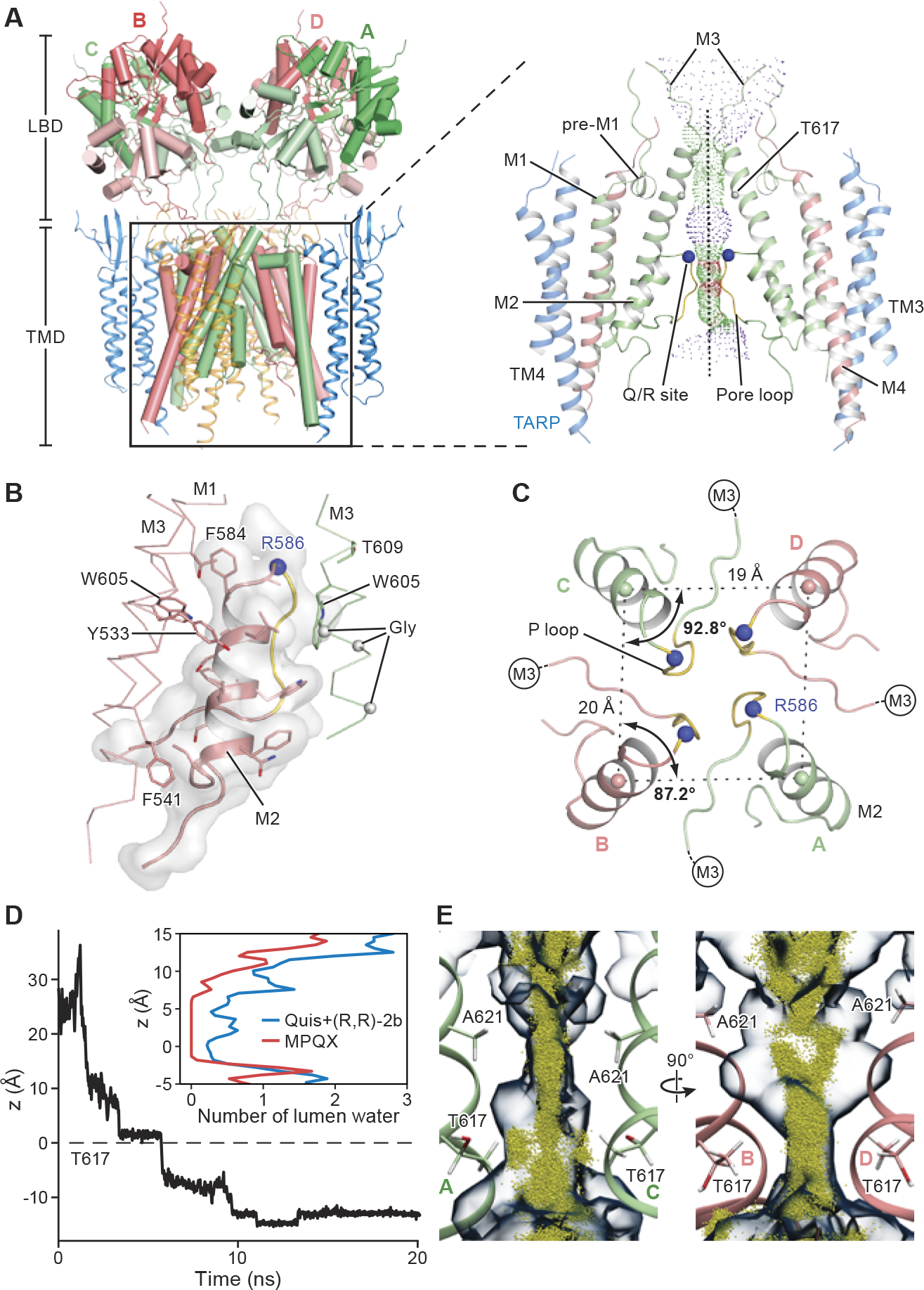
Ion channel pore of quisqualate/(R,R)-2b complex. (A) Overall view of LBD/TMD layers and, in inset, close-up view of the TMD region showing receptor TMD, TM4 of TARP and solvent accessible pathway of ion channel pore, along the central 2-fold axis (dashed line). In the inset only two subunits are shown. A/C and B/D subunits are colored in green and salmon, respectively, with the pore loops highlighted in yellow. Cα atoms of Arg586 (Q/R site) and Thr617 are shown as blue and grey spheres, respectively. Pore radii calculated without the side-chain model of Arg586 are depicted by purple, green and red dots representing pore radii of >3.3 Å, 1.8-3.3 Å, and <1.8 Å, respectively. (B) The M2 helices are stabilized through hydrophobic side-chain interactions with M1 and M3 helices from the same subunit and with M3 helix from adjacent subunits. M2 helix is represented as cartoon in transparent solvent accessible surface. Well-resolved side-chains are shown as sticks, and Cαs of key glycine residues in M3 are defined by grey spheres. (C) “Top-down” view of M2 helices and the pore loops, showing deviation from four-fold symmetry. COMs of M2 helices (salmon and green spheres), distances and angles are shown. (D) Na^+^ ion permeation trajectory captured during MD simulation of the quisqualate/(R,R)-2b structure (with R596Q mutation), showing spontaneous entry of the ion from the extracellular solution (∼20 Å away from Thr617). Inset shows the number of water molecules along the pore axis averaged over 20 ns simulation of the quisqualate/(R,R)-2b and the MPQX structures (PDB code: 3KG2). (E) Orthogonal views revealing asymmetric water and ion distributions in the channel lumen of the quisqualate/(R,R)-2b structure. The 30% water occupancy isosurface based on 80 ns SMD simulation trajectory of the quisqualate/(R,R)-2b structure is shown in semi-transparent surface. The trajectory of the permeating Na^+^ ion is shown as yellow dots.

The storied Q/R site, discovered by Seeburg and colleagues (Sommer et al., 1991b), harbors an arginine in the present construct and the four arginine residues are located at the apex of the pore loop (Figure 3A). Density for the β-carbon of Arg 586 allows us to define the orientation of the side-chain (Figure S3B), thus suggesting that the side-chains project into the central vestibule (Figure 3A), in agreement with the reduction in block by cationic toxins, small molecules and cytoplasmic polyamines (Donevan and Rogawski, 1995; Magazanik et al., 1997; Poulsen et al., 2014). The location of the arginine guanidine groups also provides a logical explanation for the calcium impermeability of GluA2 (Arg586)-containing AMPA receptors (Hollmann et al., 1991) due to charge-charge repulsion.

The M2-pore loop region of the quisqualate/(R,R)-2b complex is 2-fold symmetric (Figure 3C), in contrast with the 4-fold symmetric pore helices found in the MPQX-bound complex, suggesting that the pore structure is not independent of structural rearrangements related to complex activation. Interestingly, the pore symmetry is unaltered in potassium channels upon gate opening, perhaps because a glycine residue (Gly99 for KcsA) present in the gating helix functions as a hinge (Jiang et al., 2002), largely decoupling conformational changes associated with gating from movements of the pore helix and pore loop. The equivalent position in AMPA and kainate receptors is replaced by a threonine (Thr609 for AMPA receptors) (Figure 3B). We speculate that this renders the M3 helices more rigid, thus coupling movements of M3 helices to the structural elements of the pore via extensive M2-M3 interactions.

The M3 bundle crossing in the quisqualate/(R,R)-2b complex forms a two-fold symmetric pore that is more dilated along the B/D direction in comparison to the A/C direction, breaking the ∼4-fold symmetry observed in the antagonist-bound complex (Figures 3C and S4). To estimate whether this gate is sufficiently open to allow for ion permeation, we measured distances between Cα atoms of opposing residues including Thr617, Ala621, Thr625 and Met629 and compared them with corresponding distances measured from an inactive/closed GluA2 receptor (Sobolevsky et al., 2009). The Cα atom distances increase by 2 Å, 2 Å, 3 Å and 14 Å at Thr617, Ala621, Thr625 and Met629 between A and C subunits, respectively, and by 2 Å, 4 Å, 20 Å and 14 Å between B and D subunits. These distance increases, together with the calculation of the solvent accessible pathway along the pore axis (Smart et al., 1996) show that the pore constriction, or gate, has expanded and that Thr625 and Met629 no longer hinder ion-permeation (Figure 3A). However, the ‘gate region’ at the M3 bundle crossing is not as open as in the open state of potassium channels (Long et al., 2005), and thus we argue that the quisqualate/(R,R)-2b bound GluA2-TARP γ2 complex represents a partially open, ion conductive state.

## Ion-permeation profile of the ion channel pore

The moderate resolution of the cryo-EM density maps precludes precise placement of side-chains, thus leading to uncertainty over the dimensions and chemical character of the ion channel pore. To further address the question of whether or not the M3 bundle crossing is sufficiently open to allow for ion permeation, we performed a series of computational studies. To characterize the hydration of the pore, while taking into account thermal fluctuations of the pore-lining residues, and to investigate how opening of the gate affects this process, a series of equilibrium molecular dynamics (MD) simulations were performed on the membrane-embedded models derived from the quisqualate/(R,R)-2b bound GluA2 TARP γ2 complex, as well as from the previously reported closed structure (R586Q mutant) (Sobolevsky et al., 2009). The hydration patterns indicate that the quisqualate/(R,R)-2b bound GluA2 TARP γ2 complex can accommodate more water molecules, especially within the gate region (between z = -2 Å(Thr617) and z = 6 Å (Ala621)) (Figure 3D). The lumen of the open-gate structure is fully hydrated as measured by the water occupancy map (Figure 3E). Spontaneous entry of a Na^+^ ion into the pore lumen is observed within 10 ns in an MD simulation, further supports that the channel gate is sufficiently open to allow permeation of hydrated Na^+^ ions (Figure 3D).

## Asymmetric water/ion accessible region inside channel lumen

To gain deeper insight into the mechanism of ion permeation and ion-protein interactions, we employed steered MD (SMD) simulations to induce permeation of one Na^+^ ion through the gating region of the open-channel structure. These simulations further examine the hydration structure of the lumen in the presence of permeating ions. Interestingly, the water occupancy isosurface shows strong asymmetry along the lumen (Figure 3E). The region between the extracellular side and Ala621 is elongated towards the B/D subunits, while the region between Ala621 and Thr617 was elongated towards the A/C subunits. The region visited by the permeating Na^+^ ion (Na^+^ accessible region) exhibited the same asymmetry pattern (Figure 3E). The measured asymmetry showed that the Na^+^ accessible region near Ala621 was symmetric (Figure S5B), likely due to the hydrophobicity of its side-chain. By contrast, the Na^+^ accessible region near Thr617 was elongated (Figure S5B). Close inspection of the region near Thr617 reveals that the asymmetry is caused by favorable interactions of the Na^+^ with the individual hydroxyl groups of the lining Thr617 side-chains (Figure S5C). These interactions also lead to a rotation of the longitudinal axis of the Na^+^ accessible region by 90° at Thr617 (Figure S5D). We further note that the side-chain orientations of Thr617 from the A/C chains changed during these ion-permeation simulations, while those from the B/D chains retain their original conformation (Figure S5E), likely underlying preferential interaction between the Na^+^ ion and Thr617 from the A/C chains.

The average number of first-shell water molecules surrounding the permeating Na^+^ was 5.2, compared to 5.7 in bulk water, throughout most of the pore (region above Thr617 Cα), but dropped by 1 unit upon interacting with Thr617 (Figures S5F and S5G). A second drop in ion hydration occurred at ∼2 Å below Thr617, caused by interactions with the hydroxyl group of Thr617 from chain B (Figure 3E). This is also supported by inspecting the relation between the number of solvation shell water molecules around Na^+^ and the minimum distance between Na^+^ and the hydroxyl oxygens of Thr617 (Figure S5H), a solvation metric that decreases when Na^+^ is close to the oxygen of one of the Thr617 residues. The asymmetric distributions of water and ions within the pore were reproducibly observed in all simulations performed, despite differences in initial configurations, namely the presence of arginine or glutamine at site 586, or different protonation states of the four Arg586 residues (see Methods).

## TARP modulation of receptor gating

We observe interactions between the “KGK” motif in the LBD and the acidic loop of TARP (Dawe et al., 2016), yet only for subunits in the B/D positions in the quisqualate/(R,R)-2b and kainate/(R,R)-2b complexes (Figures 4A and 4B), emphasizing the distinct roles of TARPs in the A’/C’ or B’/D’ positions. Upon transition from the MPQX to the quisqualate/(R,R)-2b bound states, the “KGK” motif of the B/D subunits shifts by ∼7 Å yet remains near the acidic loop of the B’/D’ TARP subunits, accompanied by movement of the receptor S1-preM1 and M3-S2 linkers towards TARP (Figures 4B and 4C). The cryo-EM density suggests that the TARP acidic loop is flexible and thus we speculate that the loop could engage the “KGK” motif during the progression from antagonist-bound/inactive to agonist-bound/active states, thus providing a structural explanation for how 6-cyano-7-nitroquinoxaline-2,3-dione (CNQX) acts as an agonist on AMPA receptor – TARP assemblies (Menuz et al., 2007). By contrast, activation by quisqualate increases the COM distance between the “KGK” motif and the TARP α1 helix to a displacement that is 10 Å greater than the equivalent distance at the B/D position (Figures 4A and 4C), thus precluding interactions between the “KGK” motif and the TARP acidic loop in the A/C positions, in accord with the disorder of the A’/C’ TARP acidic loops. Instead, at the A’/C’ positions the TARP β-sheets are near the LBD S2-M4 linkers, in position to interact with the receptor flip/flop region known to modulate gating kinetics (Figure 4D) (Mosbacher et al., 1994). By contrast, these regions are distant at the B’/D’ positions (Figure 4E).

**Figure 4.**
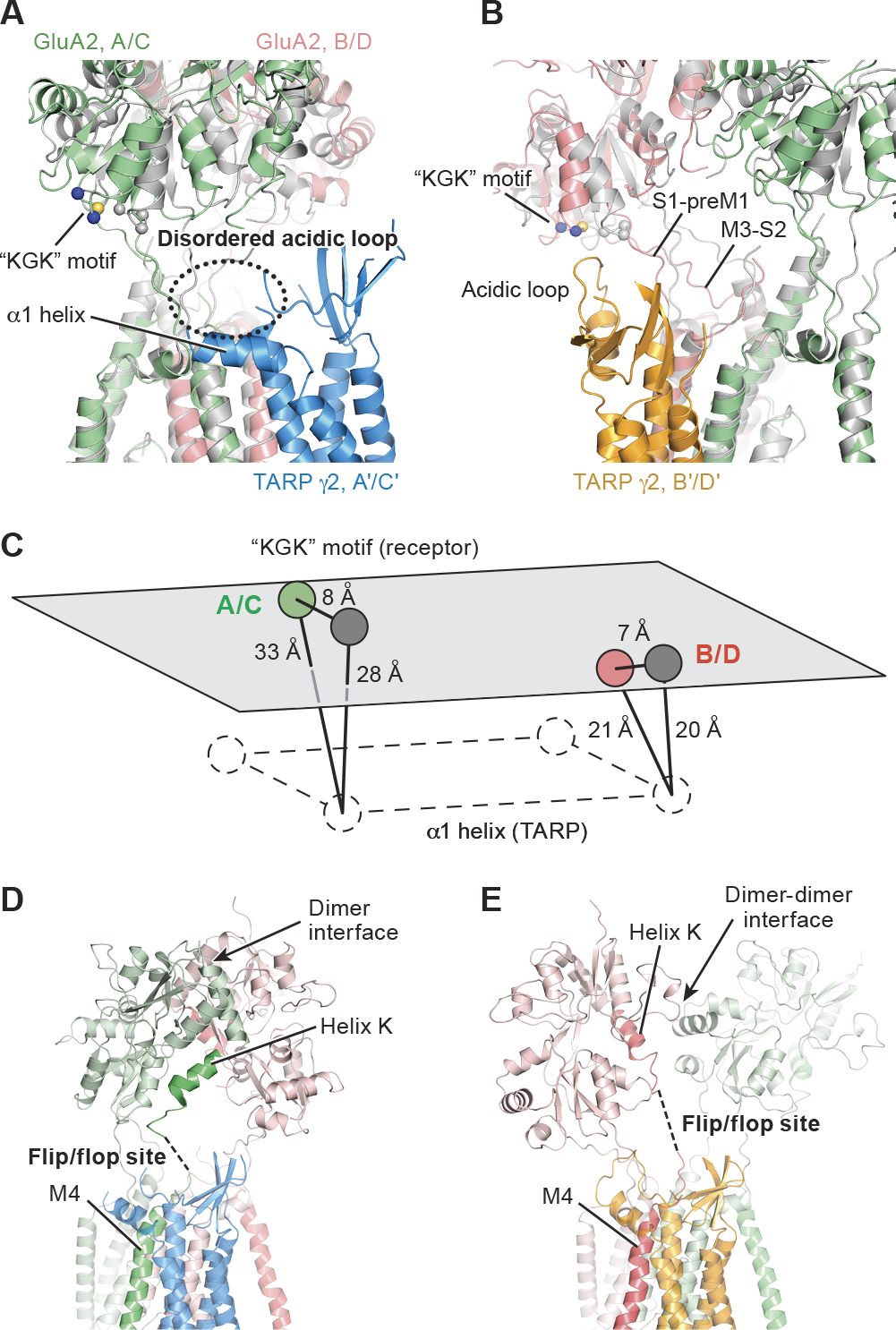
Interactions between GluA2 LBD and TARP γ2. (A) and (B) GluA2 LBD-TARP γ2 interface viewed parallel to the membrane at A/C and B/D positions, respectively. Structures of GluA2-TARP γ2 complex bound with quisqualate/(R,R)-2b (green and salmon) and MPQX (grey) were superimposed using main-chain atoms of receptor TMD. The Cα atoms of the ‘KGK’ motif (697-699) are shown as spheres. In the quisqualate/(R,R)-2b bound structure, lysine and glycine Cα atoms are blue and yellow, respectively. (C) Schematic diagram illustrating displacement of the “KGK” motif and change in distances between the “KGK” motif and TARP α1 helix upon receptor activation. (D) and (E) Possible interactions between TARP and LBD dimer interface (D) and between TARP and LBD dimer-dimer interface (E) in GluA2-TARP γ2 complex bound with quisqualate/(R,R)-2b. Unstructured S2-M4 linkers are represented by dashes lines.

## TARP restricts LBD rearrangement upon receptor desensitization

To study the mechanism of desensitization for the TARP-bound receptor, we incubated the apo GluA2-TARP γ2 with 2mM quisqualate, an agonist which profoundly stabilizes the receptor in a desensitized state (Jin et al., 2002). Examination and 2D classification of raw particles revealed, on the one hand, defined ATD and LBD layers yet on the other hand, splayed ATDs in single particles and blurred ATDs in some 2D classes (Figures 5A, 5B, S2K and S2L). Initial 3D classification yielded 5 classes with blurry structural features, presumably due to an averaging effect of intrinsic particle heterogeneity and conformational mobility of the LBD and ATD layers (Figure S6). Class 1 has iconic receptor ATD and LBD features along with 4 protrusions on the extracellular side of micelle, thus representing one conformation of the ostensibly desensitized receptor-TARP complex. Reconstructions carried out using particles from class 1 resulted in higher resolution EM map with more abundant features when C2 symmetry was imposed, compared to that without imposed symmetry (Figure S6). None of the classes had a 4-fold symmetric LBD layer, suggesting that the desensitized state(s) of the GluA2-TARP γ2 complex might be fundamentally different from kainate receptors (Meyerson et al., 2016). Although classification and reconstruction of particles in other classes confirmed the conformational heterogeneity in the LBD and ATD layers, the presence of TARP was not clearly evident for any subset of the particles obtained by various strategies. Because the AMPA receptor-TARP complex is prone to disassociation upon desensitization (Morimoto-Tomita et al., 2009), the receptor – quisqualate preparation studied here likely has receptors that are not saturated with TARP subunits. Further studies will be required to define the structures of the desensitized state of AMPA receptors not saturated by TARP subunits.

**Figure 5.**
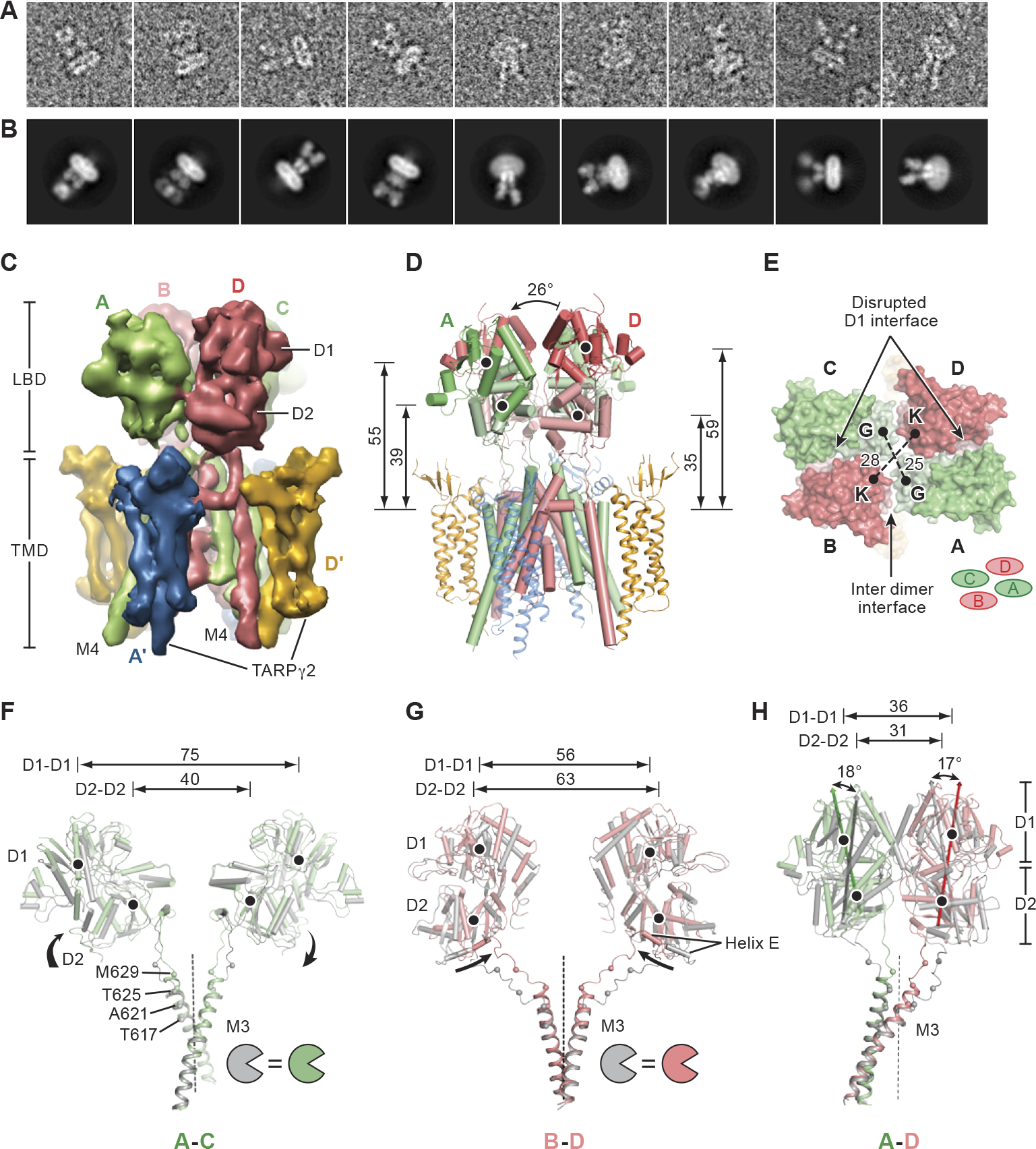
Quisqualate complex. (A) and (B) Raw particle images (A) and representative 2D-class averages (B) illustrating the conformational heterogeneity of GluA2-TARP γ2 quisqualate complex. (C) Density map of the most well defined 3D class where refinement was focused on the LBD/TMD layers. View is along the LBD dimer interface showing the separation of the D1-D1 interface and formation of the D2-D2 interface. (D) and (E) “Side” and “top-down” views of quisqualate-bound GluA2-TARP γ2 complex, showing ruptured LBD dimer interface. COM distances, measured similarly as the non-desensitized structures (Figures 1E-1G). (F)-(H) Conformational changes associated with receptor desensitization revealed by superposition of LBD-M3 pairs from quisqualate bound GluA2-TARP γ2 complex structures in the absence (in color) or presence of (R,R)-2b (in grey), using main-chain atoms of receptor TMD. Views are similar as in Figure 2. Angles formed between vectors connecting COMs of D1 and D2 lobes in complexes without and with (R,R)-2b are labeled at both A/C and B/D positions in (H). Distances are in angstroms (Å).

To further explore the class 1 conformation of the desensitized GluA2-TARP γ2 complex, we enlarged the particle set by merging class 1 with class 3, the latter of which features less prominent TARP protrusions (Figure S6). After subsequent focused classification and refinement with imposed C2 symmetry, the resolution of the obtained reconstruction was estimated at 7.7 Å (Figures 5C and S2M-S2O). Main-chain density was clearly resolved for four TARPs, the LBD layer and all receptor transmembrane helices, unambiguously guiding the model fitting. To generate an initial model we extracted LBD ‘clamshells’ from the crystal structure of the isolated LBD quisqualate complex (Jin et al., 2002) and fit each LBD into the EM density as rigid bodies; receptor transmembrane helices M1,M3 and M4 were extracted from the MPQX bound GluA2-TARP γ2 complex and also docked into the EM density as a rigid body (Zhao et al., 2016), resulting in reasonable model to map fitting. The model for the rest of the pore region and for the TARPs were derived from the quisqualate/(R,R)-2b bound GluA2-TARP γ2 complex and fitted into the EM density. All of these structural components were then refined, in real-space, against the cryo-EM map (Table S1).

The resulting structure reveals that the M3 gate is closed and the degree of LBD clamshell closure is similar to the isolated LBD structure bound with quisqualate (Figures 5D, S3D and S4), confirming that the complex is stabilized in an agonist-bound desensitized state. The ‘elevation’ of the LBD layer from the reference plane defined by Thr617 Cαs is similar to that of the agonist-bound non-desensitized states (Figure 5D). However, the gating ring has contracted upon transition from the active state to the desensitized state (Figure 5E). In addition, there is a ∼26° rotation of one subunit of a LBD dimer relative to the quisqualate/(R,R)-2b bound state (Figure 5D), rupturing the LBD D1-D1 interface while forming new D2-D2 contacts (Figures 5C, 5D and Movie S1). Strikingly, the LBD dimers adopt an approximately similar conformation as that seen in the isolated LBD complex of the GluA2 S1S2J G725C structure bound with glutamate (PDB code: 2I3W) (Armstrong et al., 2006) with a superposition of LBD dimers yielding an root mean square deviation (RMSD) of ∼1.3 Å for Cα atoms.

In contrast to the large-scale LBD rearrangement found with isolated AMPA and kainate receptors (Dürr et al., 2014; Meyerson et al., 2016; Meyerson et al., 2014), GluA2-TARP γ2 complex desensitization involves more subtle rotations of the LBD subunits, as indicated by a structural comparison between the quisqualate bound desensitized state and quisqualate/(R-R)-2b bound active state (Figures 5F-5H). In turn, the LBD E helices move towards the LBD center, releasing the mechanical pulling force exerted on the M3 helices through M3-S2 linkers (Figures 5F-5H). Given that the LBD clamshell closure in the apo state is comparable to MPQX bound state (Dürr et al., 2014; Zhao et al., 2016), the GluA2-TARP γ2 structure bound with MPQX approximates the apo state, thus allowing us to propose a mechanism of receptor resensitization. Superimposition of structures of the desensitized complex with the MPQX-bound complex reveals relatively subtle displacements in COM positions of the D1 and D2 lobes as well as the entire LBD between the two states (Figure S7 and Movie S1), demonstrating how TARPs prevent large-scale LBD rearrangement upon desensitization. We suggest that recovery from desensitization involves reformation of the D1-D1 interface following agonist unbinding and clamshell opening, without large-scale LBD movements as would be required if the LBD layer adopted a ∼4-fold symmetric structure, thus accelerating recovery relative to the non TARP-bound receptor.

## TARP modulation of AMPA receptor gating

The partial agonist kainate and full agonist quisqualate induce greater clamshell closure than the antagonist MPQX bound state and thus yield larger D2 lobe separation, correlating with the enhanced pulling force exerted on the M3 helix. Upon desensitization, D2 lobe separation returns to an antagonist-like distance by way of D1 interface disruption and LBD domain rearrangement. To describe the D2 lobe separation in these structures, we measured distances between the COMs of helix E, located on the LBD D2 lobe and directly connected to M3 through the D2-M3 linker (Figure 6A). The helix E separation within dimers increases by 9 Å and 13 Å upon transition from the MPQX state to the kainate/(R,R)-2b and quisqualate/(R,R)-2b states, respectively (Figures 6B-6D), consistent with observations in isolated LBD and receptor structures (Dürr et al., 2014; Jin et al., 2002). When comparing the COM distances between E helices from opposing subunits during the transition from MPQX to kainate/(R,R)-2b bound complex, there are no obvious changes in the A-C distance but a 9 Å increase in the B-D distance (Figures 6B and 6C). In the quisqualate/(R,R)-2b activated complex, these distances are further increased by 6 Å and 2 Å respectively (Figures 6C and 6D), yet without enlarging the ion channel pore diameter. Interestingly, the distance change between the quisqualate/(R,R)-2b and kainate/(R,R)-2b structures in the B/D direction is smaller than in the A/C direction. For the A/C subunits, the larger distance difference is the direct consequence of the different clamshell closure induced by kainate and quisqualate (Figures 2B and 2C). For the B/D subunits, however, the D2 lobe position is largely fixed by interactions with the B’/D’ TARPs (Figures S3A and S3C), even though quisqualate gives rise to greater LBD domain closure than kainate (Figures 2B and 2C). For the quisqualate-bound desensitized state, these distances are roughly comparable with the MPQX bound state (Figures 6B and 6E), thus yielding a closed ion channel gate (Figure S4).

**Figure 6.**
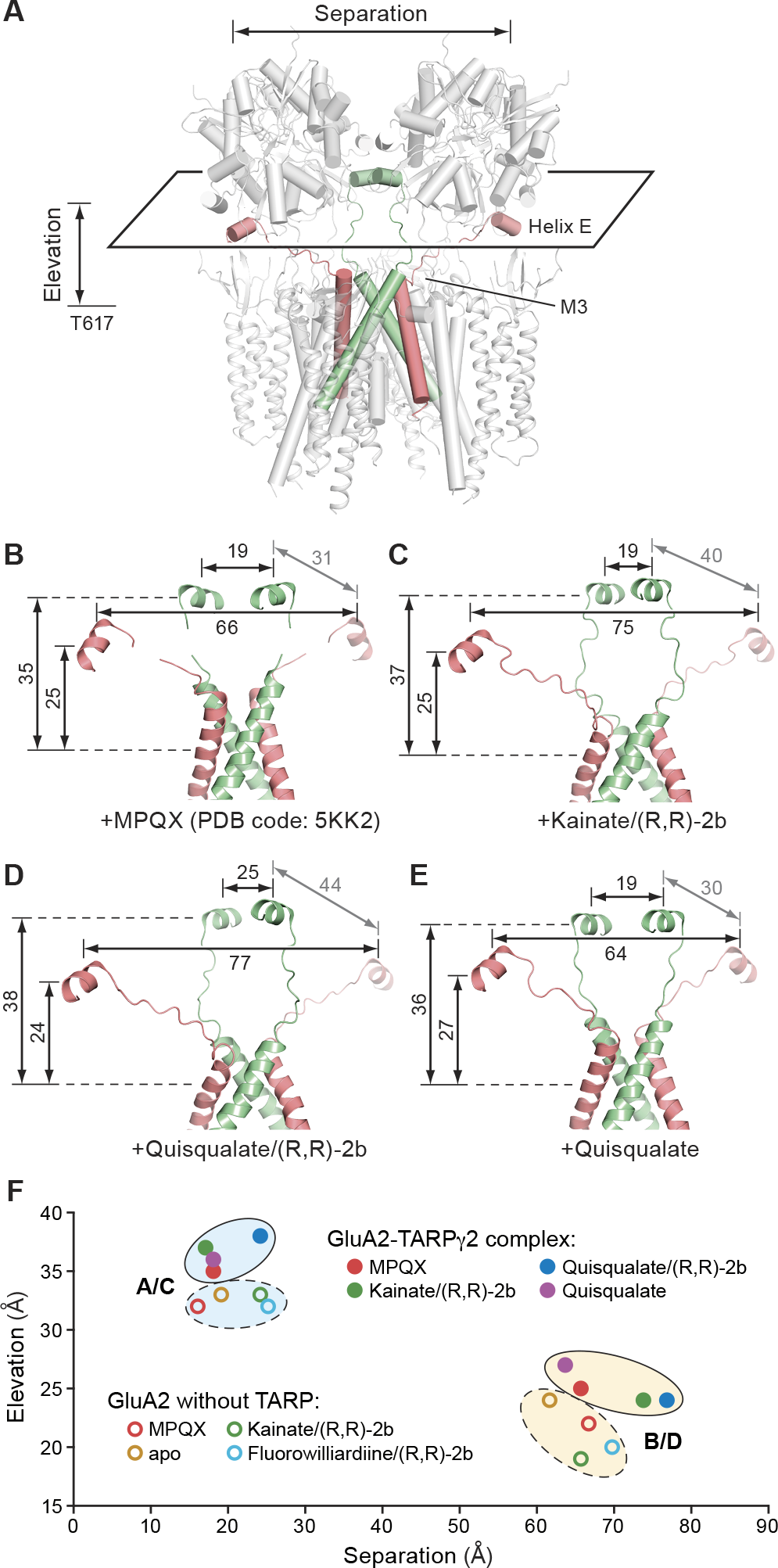
TARPs act as LBD bouy. (A) Helix E, M3 helices and the M3-S2 linker highlighted in the GluA2-TARP γ2 complex. (B)-(E) Helix E, M3 helices, and the M3-S2 linkers of GluA2-TARP γ2 complex bound with MPQX (B), kainate/(R,R)-2b (C), quisqualate/(R,R)-2b (D) and quisqualate alone (E) viewed parallel to the membrane. Distances between proximal and opposing helix E COM pairs are labeled; “elevation” is the distance perpendicular to the membrane between helix E COM and Thr617. (F) “Elevation” and “separation” plot of opposing Helix E COMs highlights the role of TARPs. Distances from the GluA2-TARP γ2 complex and intact GluA2 structures are indicated by solid and open spheres, and grouped separately in solid and dashed circles, respectively. Distances are in angstroms (Å).

Previous structural studies of the isolated GluA2 receptor demonstrated that agonist binding, in the absence of ion channel gating opening, promotes movement of the LBD layer closer to membrane (Chen et al., 2014; Dürr et al., 2014). We suggest that the movement toward the membrane releases the pulling force originated from clamshell closure and hampers channel activation. We thus measured the distances between Thr617 COM and the COM of helix E pairs in the GluA2-TARP γ2 complexes. Instead of compression to membrane, we found that helices E in the B/D subunits maintain a similar height (∼25 Å) during gating, and helices E in the A/C subunit increase in ‘elevation’ upon activation (Figures 6B-6D). In the isolated receptor structures agonist binding induces both helix E separation and compression to membrane (Figure 6F). Indeed, at equivalent states the E helices of the GluA2-TARP γ2 complex are “higher” than in the isolated receptor structure. Thus TARPs preclude LBD layer compression during gating, retaining the LBD layer in gating active state. We suggest that, in turn, tension is more efficiently exerted on the M3 helices, resulting in channel opening even with small clamshell closure, such as that induced by kainate and CNQX (Menuz et al., 2007; Tomita et al., 2005; Turetsky et al., 2005).

In this structural investigation of GluA2-TARP γ2 complex, conformational intermediates were captured by single-particle cryo-EM, elucidating a molecular mechanism of complex activation and desensitization. In the context of the GluA2-TARP γ2 complex, agonist efficacy largely determines the extent of LBD clamshell closure. When the D1-D1 interface is intact, LBD closure is transduced into expansion of D2 layer, charging the M3-S2 linkers with tension (Figure 7) and promoting opening of the ion channel gate. In the absence of TARPs, the LBD gating ring can compress to the membrane, releasing the tension required for gating opening (Figure 7). To potentiate receptors, TARPs function as a molecular buoy that prevents LBD from “sinking” towards the membrane, thus promoting the transduction of LBD clamshell closure to channel opening (Figure 7). Electrostatic interactions between the conserved “KGK” motif of the receptor and the acidic loop of TARP, at the B/D positions, allows the respective LBD clamshells to promote a larger gate opening in comparison to the A/C pair, offering a structural basis for the greater importance of B/D pair than A/C pair in channel gating (Figure 7). The “KGK” motif-acidic loop interaction increases kainate efficacy by positioning the LBD D2 lobes of the more crucial B/D pair. While the desensitized state of the GluA2-TARP complex likely involves an ensemble of structural states, a highly populated conformation shows how TARPs encircle the agonist-bound LBDs, reducing the conformational changes upon desensitization in comparison to the isolated receptor. Moreover, the LBD harbors 2-fold rather than 4-fold symmetry and exhibits only rupture of the D1-D1 interface rather than large-scale reorientation of the LBD clamshells (Figure 7). We speculate that TARPs accelerate receptor resensitization by restricting the LBD layer from large-scale rearrangements upon desensitization, thus facilitating reformation of LBD dimer interface to recreate an active GluA2-TARP γ2 complex.

**Figure 7.**
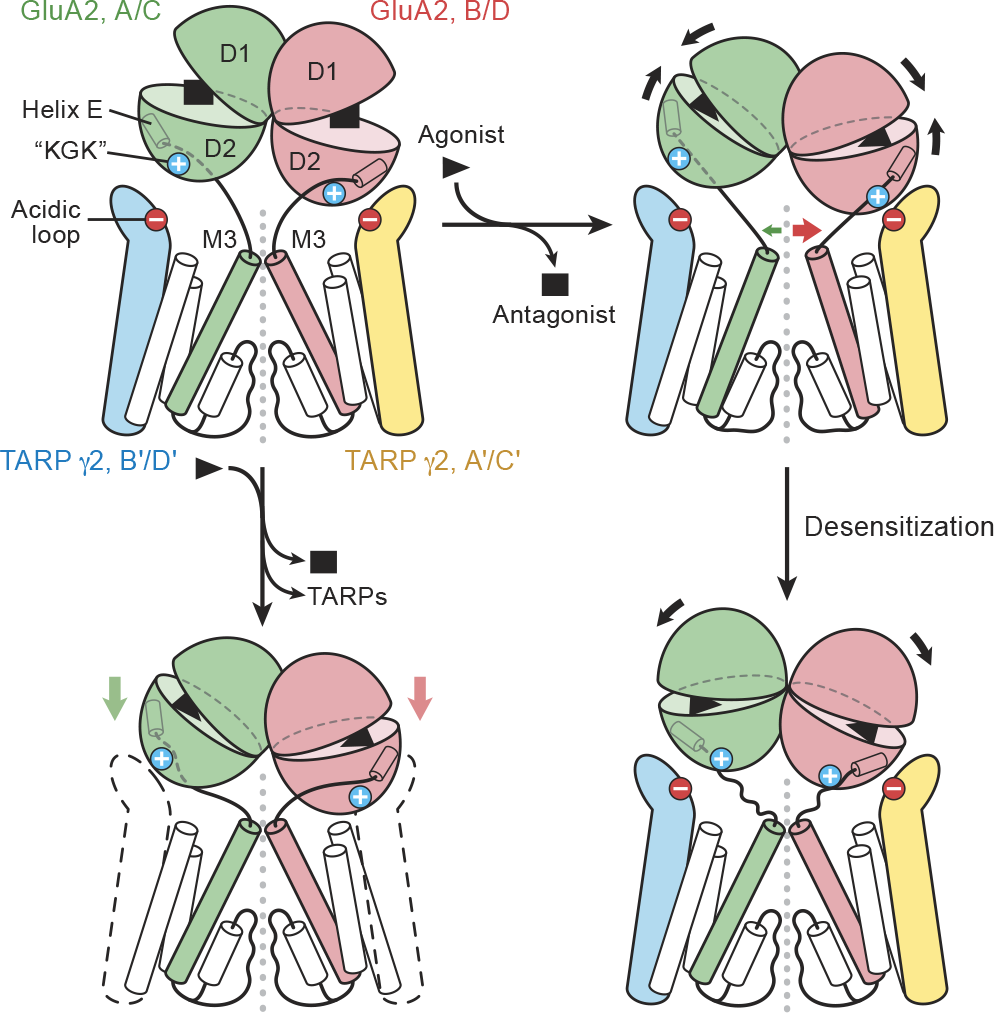
Mechanisms of receptor activation and desensitization. Shown is the LBD/TMD layer of two receptor and two TARP subunits. TARPs function as a molecular buoy on which the LBD layer ‘floats’, ensuring tension exerted on M3-S2 linkers is efficiently transmitted to open the channel gate rather than causing LBD to approach the membrane.

## Materials and methods

### GluA2-TARP γ2 expression and purification

The expression and purification of GluA2 (flop variant, arginine at the Q/R site)-TARP γ2 complex were carried out as described (Zhao et al., 2016), with minor modifications. In brief, the complex was expressed using Clone #10 cells adapted to grow in suspension (Shanks et al., 2010), cultured in Freestyle 293 expression medium supplemented with 2% (v/v) fetal bovine serum and selection antibiotics (125 μg/ml zeocin, 150 μg/ml hygromycin, and 125 μg/ml neomycin). Whereas TARP γ2 expression was constitutive, GluA2 expression was induced by addition of 7.5 μg/ml doxycycline at a cell density of 2×10^6^ cells/ml. Subsequently, 200 nM MPQX (also known as ZK200775) (Turski et al., 1998) was added to the media to prevent cytotoxicity due to receptor-TARP complex overexpression. Cells were collected by centrifugation 30∼35 hours post-induction and lysed by sonication. After removal of cell debris by centrifugation at 1,200 × g (15 min at 4 °C), the supernatant was subjected to ultracentrifugation at 100,000 × g for 1 hour to collect the membrane fraction.

The membrane fraction was resuspended and solubilized in TBS buffer (20 mM Tris, pH 8.0, 150 mM NaCl) containing 1% (w/v) digitonin and 1 μM MPQX for 2 hours at 4 °C. Insoluble material was removed by ultracentrifugation at 100,000 × g for 1 hour, and the supernatant was passed through an anti-FLAG immunoaffinity column preequilibrated with buffer P (20 mM Tris pH 8.0, 150 mM NaCl, 0.1% (w/v) digitonin), followed by a wash step using at least 20 column volumes of buffer P to remove contaminants and MPQX. The FLAG-tagged GluA2 receptor in complex with TARP was eluted with buffer P supplemented with 0.5 mg/ml FLAG peptide. For samples with positive allosteric modulator, the eluted complex was concentrated and further purified by size-exclusion chromatography (SEC) using a Superose 6 10/300 GL column equilibrated in Buffer P supplemented with 1 μM (R,R)-2b. Peak fractions were combined, supplemented with additional (R,R)-2b to a final concentration of 50 μM and concentrated to 3 mg/ml using 100-kDa cutoff concentrator. For studies of in the absence of modulator, SEC and protein concentration were carried out following the same procedure without (R,R)-2b.

### Cryo-EM data acquisition

A droplet of 2.5 μl of purified GluA2-TARP γ2 complex at 3 mg/ml was deposited on Quantifoil 1.2-1.3 Au 300 mesh grids glow discharged at 15 mA for 120 s. The grid was then blotted for 3.5-4.5 s at 22 °C under conditions of 100% humidity, and flash-frozen in liquid ethane.

Cryo-EM data were collected on a 300 kV microscope using a direct electron-detection camera (K2 Summit) positioned post a GIF quantum energy filter. The energy filter slit width was set to 20 eV and a 100 μm objective aperture was used. Micrographs were recorded in super-resolution counting mode at a magnified physical pixel size of 1.72 Å, with the defocus values ranging from 1.3 to 2.2 μm. A 20 s exposure was fractionated into 50 frames, each exposed for 0.4 s at a dose rate of 8.0 e^-^ /pix/s, resulting in a total dose of 54 e^-^/Å^2^.

### Image processing

For the quisqualate/(R,R)-2b bound complex, a total of 4470 micrographs were corrected for beam-induced drift using UCSF MOTIONCORR2 (Zheng et al., 2016). The contrast transfer function (CTF) parameters for each micrograph were determined by CTFFIND4 (Rohou and Grigorieff, 2015) and particles were picked using DoG-picker (Voss et al., 2009) to minimize template bias. Contrast-based particle-picking generated a particle set contaminated by crystal ice, micelles, disassociated or disordered protein and other false positives, which were largely removed by rounds of 2D classification, resulting in classes with recognizable features in agreement with 2D projections calculated from the crystal structure of the antagonist-bound receptor (PDB code: 3KG2) (Sobolevsky et al., 2009). From an initial set of 726.6k putative particles, 236.7k particles were selected for subsequent 3D classification.

This subset of particles was classified into six 3D-classes using a reference model generated from the cryo-EM map of the MPQX-bound GluA2-TARP γ2 complex, which had been low-pass filtered to 40 Å (Zhao et al., 2016). Four of the resulting classes, featuring ∼4-fold symmetric protrusions on the extracellular side of the detergent micelle, were combined to yield a 160.5k particle stack. Using a soft mask extending ∼8 Å from the LBD and TMD domains with an additional ∼5 Å cosine edge, along with C2 symmetry, refinement in Relion resulted in a reconstruction at 5.5 Å resolution as estimated by Fourier shell correlation (FSC) between two independently refined half-maps, using the 0.143 cutoff (Scheres, 2012). The particle CTF parameters were locally refined using Gctf (Zhang, 2016), which improved density map features and the FSC resolution to 5.4 Å. Finally, aligned particles were subjected to a 3D-classification focused on LBDs and M3 helices without further particle alignment. The most populated class, containing 144.2k particles, was used to repeat refinement, producing the final reconstruction at 4.9 Å resolution. To access the quality of data and reconstruction, FSC between two independently refined half-maps and angular distribution of particles used for refinement were plotted, and local-resolution throughout the map was calculated using ResMap (Kucukelbir et al., 2014).

Image processing and reconstruction for the kainate/(R,R)-2b bound complex was carried out similarly, starting from 2374 micrographs, to a final particle set containing 84.5k particles. Relion-based refinement resulted in a 6.4 Å reconstruction. For the quisqualate-bound GluA2-TARP γ2 complex, four batches of data consisting of 4275 micrographs in total were drift-corrected by Unblur (Grant and Grigorieff, 2015). CTF parameters were estimated by CTFFIND4 (Rohou and Grigorieff, 2015) and particles were picked by Gautomatch (www.mrc-lmb.cam.ac.uk/kzhang/Gautomatch/). In comparison to micrographs of particles with allosteric modulator (R,R)-2b, the particles without (R,R)-2b were more sparsely distributed on the micrographs. Moreover, the fraction of particles remaining after removal of false positives by 2D-classification was substantially lower, possibly because the receptor-TARP complex is less stable in the agonist-bound/desensitized state. As a result, only 134.8k particles were subjected to 3D-classification, yielding 5 classes. Classes 1 and 3 featured four protrusions at the extracellular side of micelles, contained ∼50% of the total particles, and clearly represent the GluA2-TARP γ2 complex, although the TARP protrusions in class 1 are more prominent than in the third class. Therefore, the initial 3D reconstruction was carried out with the particle set from the first class using a soft mask containing LBD and TMD layers and with either C1 or C2 symmetry. The reconstruction with C2 symmetry yielded a ∼8.2 Å map featuring more continuous transmembrane helices than the reconstruction with C1 symmetry and the LBD domains were also better defined in the C2 map in comparison to the C1 symmetry map, thus justifying the imposition of C2 symmetry. Even though class 3 was possibly heterogeneous in TARP occupancy, as suggested by weak TARP features, we wanted to employ as many fully occupied complex particles as possible and thus classes 1 and 3 (64.4k particles in total) were combined for an additional round of 3D classification in Relion with C2 symmetry and the LBD-TMD soft mask. This classification gave rise to a class containing 47.9% of total input particles, which was thereafter subjected to 3D refinement focused on the LBD and TMD layers with C2 symmetry. The final reconstruction of the desensitized GluA2-TARP γ2 complex bound with quisqualate was determined at 7.7 Å based on FSC and substantiated by the stereochemical quality of the density features.

### Structural modeling

The structural modeling for the quisqualate/(R,R)-2b bound GluA2-TARP γ2 complex commenced by rigid-body fitting of D1 (residues 391-497, 731-774) and D2 (residues 498-505, 633-730) lobes extracted from the crystal structure of isolated LBD bound with quisqualate (PDB code: 1MM6) (Jin et al., 2002) and receptor TMD and TARPs extracted from the MPQX-bound GluA2-TARP γ2 cryo-EM structure (PDB code: 5KK2) (Zhao et al., 2016) into the cryo-EM density. The density map for receptor and TARP TMDs are rich in features including grooves for helices and side-chain density for most aromatic residues, ensuring an accurate register assignment. In particular, we improved the model for the M2 pore helices and the adjacent loops (residue 566-595) following well-resolved continuous density. The ResMap estimated local resolution of this region was higher than the overall resolution, consistent with clearly visible side-chain density for Leu577, Trp578, Phe579, Leu581 and Phe584, thus providing a reliable structure for the M2 pore helix (residues 574-585). Next, the receptor TMD model was subjected to molecular dynamics flexible fitting (MDFF) against the density map (see below) (Trabuco et al., 2008), which improved the correlation coefficient between model and map from 0.58 to 0.62. Subsequent manual adjustments including fitting the backbone of S2-M3 loops (residue 625-632) into the density, deletion of un-resolved side-chains and subtle local alteration for stereochemistry optimization and clash minimization were carried out in COOT (Emsley and Cowtan, 2004).

Structural modeling for the GluA2-TARP γ2 complex in the presence of kainate and (R,R)-2b was achieved by first fitting the D1 and D2 lobe from the crystal structure of the kainate and (R,R)-2b bound LBD (PDB code: 4U1O) (Dürr et al., 2014) into the density map as rigid bodies. The TMD model was obtained by extracting receptor and TARP TMDs from the structure of GluA2-TARP γ2 complex bound with quisqualate and (R,R)-2b and fitting them into the 6.4 Å density map, resulting in satisfying map-model fitting. Residues near the hinge and LBD-TMD linkers were subject to local adjustment to optimize the stereochemistry in COOT (Emsley and Cowtan, 2004).

Structural modeling for the GluA2-TARP γ2 complex in the presence of quisqualate and absence of (R,R)-2b was carried out by rigid body fitting. A protomer was extracted from the crystal structure of GluA2 LBD bound with quisqualate (PDB code: 1MM6) (Jin et al., 2002) fit into the density for each of the four receptor LBDs. TARP and receptor pore structure (M2 helix and pore-loop) were extracted from the cryo-EM structure of GluA2-TARP γ2 complex bound with quisqualate and (R,R)- 2b. A model for the preM1-M1, M3 and M4 was extracted from MPQX bound cryo-EM structure of GluA2-TARP γ2 complex (PDB code: 5KK2) (Zhao et al., 2016). These model components were docked into the density map separately, each as a single rigid-body.

All of three initial models were further refined in real space by Phenix against the corresponding cryo-EM map, with secondary structure, 2-fold non-crystallographic symmetry and Ramachandran restraints applied throughout the refinement (Adams et al., 2010). The correlation coefficient between the refined model and map around atoms present in the model was improved from 0.62 to 0.70 in quisqualate/(R,R)-2b bound state, from 0.80 to 0.83 in the kainate/(R,R)-2b bound state and from 0.69 to 0.72 in quisqualate-alone state, indicating reasonable model-map agreement. The refined model was also converted into a density map to calculate the Fourier shell correlation with the experimental density map. The model stereochemistry was evaluated using MolProbity (Chen et al., 2010).

### MDFF structure refinement

Prior to the MD studies, MDFF was used as part of the structure refinement process (Chan et al., 2011; Singharoy et al., 2016; Trabuco et al., 2008). During the MDFF refinements, generalized Born implicit solvent model (Still et al., 1990) was used with a 0.15 M ionic strength. A 1-fs time step was used with van der Waals interactions evaluated every 2 fs and electrostatic interactions every 4 fs. Two-fold symmetry restraints were applied to the protein Cα atoms with a spring constant of 1 kcal/mol/Å^2^/atom (Chan et al., 2011). Chirality and secondary structure restraints were applied to the protein. A density map derived grid-force potential was applied to protein heavy atoms. Each heavy atom was assigned with a virtual charge of +1 and a scaling factor equal to its atomic mass. The structure was energy-minimized for 400 steps and simulated for 80 ps.

### MD simulation setup

Three types of MD simulations were performed: equilibrium MD, steered MD (SMD) (Izrailev et al., 1999; Lu and Schulten, 1999), and confined MD (CMD). In SMD simulations, the structure reported in this study excluding the LBD was used as the initial structure. Side-chain conformations were optimized using SCWRL4 (Wang et al., 2008). Residue E191 from the TARP was protonated based on pKa estimation using PROPKA 3.1 (Olsson et al., 2011). Residue 586 (Q/R site (Sommer et al., 1991a)) can be either Q or R, depending on RNA editing, and both variants were simulated in SMD. Simulation of the Q variant is denoted as “Q” hereafter. Due to the ambiguity of the protonation state of Arg586, we explored all possible variants of protonation states of the four side-chains, resulting in different net charges: 0 (R0), +1 (R1), +2 (R2a and R2b, referring to neighboring and diagonal positioning, respectively), +3 (R3) and +4 (R4).

In all simulations, the protein was first embedded in a pure POPC lipid bilayer and solvated with TIP3P (Jorgensen et al., 1983) water and 0.15 M NaCl. The systems (∼244,000 atoms, 160 Å × 160 Å × 110 Å) were then energy-minimized for 500 steps and simulated for 0.5 ns at 310 K with all protein heavy atoms and lipid phosphorus atoms harmonically restrained (k = 5 kcal/mol/Å^2^) to allow for relaxation of lipid tails. This step was then followed by a 4-ns membrane relaxation, only restraining the Cα atoms from the well-structured (helix/β-sheet) regions (k = 50 kcal/mol/Å^2^). An electric potential of -100 mV (negative at the cytoplasmic site) was added to all simulations to represent the membrane potential. In SMD simulations, starting from an equilibrated system, one Na^+^ ion was placed near Ala621 at the beginning of the simulation and was steered towards the cytoplasmic side along the Z axis (membrane normal) at a constant velocity of 0.2 Å/ns and using a spring with k = 20 kcal/mol/Å^2^ for 40 ns. No restraining forces were applied in the XY plane. An additional SMD simulation (84.5 ns) of the Q variant (without LBD/TARP) was also performed to explore the entire pore region between the extracellular side and the channel central cavity, where an ion was initially placed 4 Å above Ala621 and followed the same SMD protocol stated above.

To gain enhanced sampling of Na^+^-Thr617 interactions, CMD simulations were also performed in which two harmonic potentials were used to sandwich the ion and confine its diffusion to the region near Thr617. Using NAMD grid forces (Wells et al., 2007), two half harmonic potentials (k = 50 kcal/mol/Å^2^) were added, one near Thr617 (3 Å below its Cα) and another 4 Å above it, respectively. Two Q/R site variants (Q and R0) were simulated for 60 ns each. To test the ion permeability, equilibrium MD simulation of the Q variant was performed for 89 ns in the presence of 500 mM NaCl with -300 mV membrane potential following the same system preparation protocol.

In addition, to measure the hydration profile of the channel lumen, equilibrium MD simulations of the closed structure (PDB ID: 3KG2) and the open-gate structure reported here were performed for 24.5 ns each. Missing loop regions in the closed structure were modeled using MODELLER (Fiser et al., 2000; Sali and Blundell, 1994; Shen and Sali, 2006; Webb and Sali, 2014). Furthermore, a short MD simulation (100 steps of energy minimization and 5 ns equilibration) of one Na^+^ and one Cl^-^ ion in bulk water (50 Å × 50 Å × 50 Å) was also performed as a control to quantify the hydration of an isolated Na^+^ ion. The last 4 ns of this simulation was used for radial distribution function calculation of Na^+^ and the oxygen of water as well as the average number of water in the first hydration shell of Na^+^.

### Simulation protocol

All simulations were performed with NAMD2 (Kalé et al., 1999; Nelson et al., 1996; Phillips et al., 2005) using the CHARMM36 force field (Huang and MacKerell, 2013) under the NPT ensemble with periodic boundary conditions. The simulation temperature was controlled by Langevin dynamics with a damping coefficient of 5 ps^-1^. The pressure of the system was kept at 101.325 kPa using the Nosé-Hoover Langevin method (Feller et al., 1995; Martyna et al., 1994) with a piston period of 200 fs and piston oscillation decay time of 100 fs. Long range electrostatic interactions were calculated using the particle mesh Ewald method (Darden et al., 1993; Essmann et al., 1995) with a maximum grid spacing of 1 Å. 2-fs time step was used with short-range nonbonded interactions calculated every 2 fs and long range electrostatics every 4 fs. Conjugated gradient algorithm was used for energy minimization.

### Analysis of Na^+^-accessible region

The asymmetry in the shape of the region inside lumen available to the Na^+^ ion (Na^+^-accessible region) was defined as the ratio between the standard deviation of the Na^+^ trajectory along the first and second principal axis during the SMD simulation. First, principal component analysis (PCA) of the xy plane projection of the first 2-ns trajectory (10,000 data points) of the steered Na^+^ ion was performed. The first and second eigenvectors (sorted by decreasing eigenvalues) were chosen as the first and second principal axes, respectively. Then the coordinate system of the data points was changed to the one defined by the two principal axes by projection transformation, which effectively aligns the Na^+^ trajectory so that the direction with the largest deviation for the center is aligned with the x axis. After the projection transformation, the asymmetry defined by the ratio between the standard deviations along x axis and y axis was calculated. The corresponding z coordinate was taken from the average z positions of this 2-ns trajectory. Like the running average calculation, the asymmetry at other z locations was calculated by sliding a subsampling window (10,000 data points) along the entire trajectory. The orientation of the Na^+^-accessible region was defined as the angle between the first principal axis and the x-axis from the coordinate system of the simulation and calculated using the sliding subsampling windows stated above.

All the figures were prepared with Pymol, UCSF Chimera (Pettersen et al., 2004), VMD (Humphrey et al., 1996) and Prism 5.

## ACKNOWLEDGEMENTS

We thank T. Nakagawa for providing the clone #10 cell line, the Multiscale Microscopy Core (OHSU) for support with microscopy, and the Advanced Computing Center (OHSU) for computational support. We are grateful to L. Vaskalis for assistance with figures, H. Owen for help with proofreading and other Gouaux laboratory members for helpful discussions. S.C. is supported by an American Heart Association postdoctoral fellowship (16POST27790099). This work was supported by the NIH (NS-038631 to E.G., and P41-GM104601 and U54-GM087519 to E.T.). Computational resources were provided by XSEDE (MCA06N060 to E.T.). E.G. is an investigator with the Howard Hughes Medical Institute.

## AUTHOR CONTRIBUTIONS

S.C., Y.Z. and E.G designed the project. S.C. and Y.Z. performed sample preparation and cryo-EM data collection. The quisqualate/(R,R)-2b reconstruction and the kainate/(R,R)-2b quisqualate-alone reconstructions were carried out by S.C. and Y.Z., respectively. S.C. and Y.Z. built, refined and analyzed the models. Y.W., M.S. and E.T. designed the computational component of the project. Y.W. and M.S. performed and analyzed the MD simulations. S.C., Y.Z., Y.W., M.S., E.T., and E.G. wrote the manuscript.

## AUTHOR INFORMATION

The three-dimensional cryo-EM density maps of the GluA2-TARP γ2 complex in quisqualate/(R,R)-2b bound state, kainate/(R,R)-2b bound state and quisqualate bound state have been deposited in the EM Database under the accession codes EMD-8721, EMD-8722 and EMD-8723, respectively, and the coordinates for the structures have been deposited in Protein Data Bank under accession codes 5VOT, 5VOU and 5VOV, respectively. The authors declare no competing financial interests. Correspondence and requests for material should be addressed to E.G.

---

**Figure S1. Work-flow of cryo-EM data processing for the quisqualate/(R,R)-2b complex.** A total of 726.6 particles were picked using DoG-picker from 4470 motion-corrected micrographs. After removing false positives including crystal ice, detergent micelles and disassociated or disordered protein by rounds of 2D classification, 236.7k particles were selected for 3D classification using 40 Å low pass filtered MPQX complex map as a reference. Four of the six classes have clear features extruding from the extracellular side of the micelle and share similar an overall shape, indicative of receptor-TARP complex of reasonable conformational homogeneity. These classes were merged and subjected to 3D refinement using a soft mask focused on LBD/TMD domains, with C2 symmetry imposed. The resulting map reaches 5.5 Å resolution as estimated by gold-standard FSC criteria. Next, the local CTF parameters of each particle were estimated by Gctf, which improved the map quality and FSC resolution to 5.4 Å. Finally, aligned particles were subject to 3D classification focused on the M3 helices and the LBD layer, giving rising to one major class containing 144.2k particles. This particle set was further refined to yield a reconstruction of 4.9 Å resolution and a density map with generally well defined main-chain for the LBD/TMD layer and with abundant side-chain features. An analogous work flow was carried out for kainate/(R,R)-2b complex (see below).

**Figure S2. Cryo-EM analysis of GluA2-TARP γ^2^ complex in different conformational states.** (A-E) Quisqualate/(R,R)-2b bound non-desensitized state; (F-J) Kainate/(R,R)-2b bound non-desensitized state; (K-O) quisqualate bound desensitized state. (A), (F) and (K) A representative electron micrograph. Several particles in side views are marked by white circles. (B), (G) and (L) Selected two-dimensional class averages. (C), (H) and (M) FSC curves calculated between two independently refined half-maps before (red) and after (blue) post-processing, overlaid with FSC curve calculated between cryo-EM density map and structural model shown in grey. (D), (I) and (N) Angular distribution of particles used in the final reconstruction. (E), (J) and (O) The three-dimensional map is colored according to local resolution estimation.

**Figure S3. Cryo-EM maps and structural models for agonist-bound, non-desensitized GluA2-TARP γ^2^ complex in agreement.** (A) Dissected views of overlaid cryo-EM map and structural model of quisqualate/(R,R)-2b complex, revealing A/C and B/D positions separately. Maps and models are shown as in transparent surface and cartoon representations, respectively. (B) EM density of B’/D’ TARP subunits and each transmembrane helix of the B/D subunits of the receptor derived from the quisqualate/(R,R)-2b complex. (C) Dissected views of overlaid cryo-EM map and structural model of kainate/(R,R)-2b complex, revealing A/C and B/D positions separately. (D) Superposition of B/D LBD models in present complexes with corresponding crystal structures of isolated LBD determined in the presence of the same ligand.

**Figure S4. Cryo-EM density maps of the GluA2-TARP γ^2^ in non desensitized and desensitized states showing asymmetrical gate dilation.** (A) Overall cryo-EM maps for GluA2-TARP γ^2^ complex bound with quisqualate -(R,R)-2b, kainate -(R,R)-2b and quisqualate. (B) Cross-sections of cryo-EM map of TARP-LBD interface layer in different conformational states at positions indicated in (A)

**Figure S5. Pore structure, hydration and ion permeation.** (A) Transmembrane helices comparison between GluA2-TARP γ^2^ complex structure bound with quisqualate – (R,R) -2b and isolated receptor structure bound with fluorowillardiine -(R,R)-2b. All of the helices are shown as cylinder and the isolated receptor is colored in grey. (B)-(H) Results from the SMD simulation of the quisqualate/(R,R)-2b structure (with R586Q mutation). (B) Asymmetry of Na^+^ accessible region with the Thr617 and Ala621 regions highlighted in yellow. (C) Na^+^ ion distribution near Thr617, shown in top view (left) and two side views (right). (D) Change in orientation of the Na^+^ accessible region. (E) Trajectories of the z-coordinates of the permeating Na^+^ (black) and the hydroxyl oxygen atoms of the four Thr617 residues (colored). (F) Radial distribution function between the permeating Na^+^ ion and all water oxygen atoms calculated based on a 4-ns MD simulation of Na^+^ and Cl-in bulk water. The boundary of the first solvation shell (3.15 Å; marked with blue dotted line) was used for solvation shell analysis. (G) Variation in the number of water molecules in the first hydration shell of permeating Na^+^ ion along the channel pore axis (z-axis). (H) Relationship between the minimum distance of Na^+^ from Thr617 hydroxyl oxygen atoms and the water count in the solvation shell. The z-coordinates in panels B, D, E, and G were centered at Thr617 Cα. Raw data is shown as semitransparent lines while the running average is shown as solid lines.

**Figure S6. Work-flow of cryo-EM data processing of GluA2-TARP γ^2^ quisqualate complex.** A total of 747k particles were picked using Gautomatch from 4250 motion-corrected micrographs. After removing the false positives including crystal ice, detergent micelles and disassociated or disordered protein by multiple rounds of 2D classification, 134.8k particles were subjected to 3D classification, yielding 5 classes. Two classes feature the typical shape of full-length AMPA receptor and ∼4-fold related extrusions on the extracellular side of detergent micelle, thus putatively populated with receptor-TARP complex. The first class is clearly richer in structural features and was thereafter subjected to the initial reconstructions focused on LBD-TMD layers, with C1 and C2 symmetry, separately. The reconstruction carried out with C2 symmetry yielded a map with more features than that obtained with C1 symmetry, justifying the presence of C2 symmetry in this subset of particles. To further exploit particles adopting C2 symmetry, the two classes were combined before subjected to 3D classification focused on LBD and TMD layers, also with C2 symmetry imposed. The most populated class, containing 31.8k particles shows prominent TARP features, and was subsequently subjected to 3D reconstruction with a soft mask containing LBD and TMD in use and C2 symmetry imposed. The resolution of the final reconstruction was estimated to be 7.7 Å by FSC.

**Figure S7. Structural rearrangements necessary for receptor resensitization.** Using main-chain atoms of receptor as a reference, structural comparison was carried out between GluA2-TARP γ^2^ structures bound with quisqualate (in color) and MPQX (in grey) focusing on LBD-M3 from opposing subunits, A-C (A) and B-D (B), and adjacent subunit A-D (C). The COMs of D1 lobe, D2 lobe and entire LBD are marked by spheres.

**Movie S1. Conformational changes in LBDs and M3 helices throughout GluA2-TARP γ^2^ complex gating cycle illustrated by morphing from antagonist-inhibited, via partial and full agonist activated, to full agonist-bound, desensitized state.**

